# Plant species within Streptanthoid Complex associate with distinct microbial communities that shift to be more similar under drought

**DOI:** 10.1101/2022.08.05.502885

**Authors:** Allie N. Igwe, Ian S. Pearse, Jessica M. Aguilar, Sharon Y. Strauss, Rachel L. Vannette

## Abstract

Prolonged water stress can reduce plant growth and fitness and may shift rhizoplane microbial communities. Although drought affects microbial structure of many plant species, it is still poorly understood whether closely related plant species host distinct microbial communities or respond similarly to drought. To explore this question, eight members of the *Streptanthus* clade with varying affinity to serpentine were subjected to three watering regimes. Rhizoplane bacterial communities were described using 16S rRNA gene amplicon sequencing and we explored the impact of watering treatment and plant species on alpha diversity, differential abundance, and beta diversity of soil bacterial communities. Among most plant species, reduced watering also reduced alpha diversity, but plant species did not strongly impact alpha diversity. Drought treatment reduced microbial community dissimilarity among samples: bacterial communities were more similar when plants received less water. Watering altered the relative abundance of bacterial genera within Proteobacteria, Firmicutes, Bacteroidetes, Planctomycetes, and Acidobacteria, which responded similarly in the rhizoplane of most plant species. These results suggest that prolonged water stress in *Streptanthus* spp. can result in convergent microbial communities among plant species. We suggest the functional consequences of these shifts be examined to assess effects on plant and microbial fitness under drought conditions.

## INTRODUCTION

Many plant species have evolved physiological adaptations to withstand drought, including shorter heights, smaller leaves, and fewer stomata (Brady et al. 2005; Grant 2012; Heschel et al. 2017). In addition to, and sometimes in lieu of those traits, microbial associations can alter plant phenotype and the environmental range of conditions plants are able to tolerate (Lau and Lennon 2012; David et al. 2019). Furthermore, plants can enrich the abundance of drought-tolerant microbes or alter microbial composition in the rhizosphere which can protect plants from drought by maintaining essential functions under stressful conditions (Xu and Coleman-Derr 2019). For example, Gram-positive bacteria are considered more drought tolerant than Gram-negative bacteria and are often enriched under drought conditions (Chodak et al. 2015; Chen et al. 2020). In particular, many endospore-forming Firmicutes and Actinobacteria have been shown to increase in abundance under water stress (Wang et al. 2020). Plant-microbe interactions can influence plant reproduction, distribution and biodiversity; therefore, understanding how plants and their rhizoplane microbial communities respond to and tolerate extreme climate events, such as drought, is increasingly important for maintenance of plant diversity, production and ecosystem functions across systems undergoing rapid changes in water regimes (Cook et al. 2018).

Serpentine soil is a naturally stressful soil environment with low essential plant nutrients and low-water-holding capacity, making it drought prone (Safford et al. 2005). These soil properties pre-expose rhizosphere and the more closely root-associated rhizoplane microbial communities to stressful conditions that may elevate the abundance of drought-tolerant taxa compared to the rhizoplane microbial communities in non-serpentine soils. Under periods of water stress, drought-tolerant microbial community members from serpentine soil may proliferate and protect the plant from future drought events (Bouskill et al. 2013; de Oliveira et al. 2020). Selection of drought-resistant microbes through successive environmental filtering is one way that root-associated microorganisms from drought-prone soils may enhance plant tolerance to abiotic stress. Still, it is the interaction between abiotic conditions, plant selection, and source microbes that shape rhizoplane microbial communities.

The microbial community associated with plant roots is affected both by the plant species and also by the abiotic environment in which the plant grows. Closely related plant species can be strongly affected by underlying soil characteristics and still have distinct microbial communities when grown in a common soil environment (Burns et al. 2015; Erlandson et al. 2018). Plant species that can tolerate naturally drought-prone soils may also be uniquely able to resist perturbation by drought by leveraging associations with soil microorganisms (Brady et al. 2005). Experiments examining the relative contributions of soil type and species on structuring rhizoplane microbial communities will yield insights into how rhizoplane microbial communities of closely related species may be impacted by the same stress event (Brady et al. 2005; Naylor et al. 2017).

Here, we assess the effect of drought stress on rhizoplane microbial community composition of eight wildflower plant species within the *Streptanthus* clade. Using 16S rRNA amplicon sequencing we aimed to answer the following questions: (1) How do watering treatment, plant species, and soil affinity of the plant species impact the alpha diversity of bacterial phyla? (2) How does watering treatment shift community composition of rhizoplane bacteria? (3) Does plant affinity to serpentine soil predict rhizoplane community changes with watering treatment?

## METHODS

### Study system

The “Streptanthoid Complex” is a clade composed of several genera within the *Brassicaceae* family including *Streptanthus* spp. and *Caulanthus* spp. (Pepper and Norwood 2001; Burrell et al. 2011). Most members of this complex are annual plants that occupy rocky outcrops and slopes (Al-Shehbaz and Mayer 2008). Their presence on these bare habitats are generally a precursor to endemism, which is relatively common within the clade (Armbruster 2014; Cacho and Strauss 2014; Cacho et al. 2014). This clade is composed of ∼40 species with a range of edaphic specialization (i.e., serpentine tolerant and intolerant) (Burrell et al. 2011), making it an excellent system to study rhizoplane microbial community response to water stress. We chose eight species (or populations within a species) within this clade, four with an affinity for serpentine soils and four without an affinity for serpentine soils. (Table 1; The Calflora Database 2016). Serpentine affinity varies among species in this clade, from serpentine specialists to species with populations that can grow across a large range of substrates, including serpentine.

**Table 1.** Species names, serpentine affinity, and natural rainfall. Rainfall data obtained from The Calflora Database. Serpentine affinity describes the likelihood of finding a particular plant on serpentine soil from serpentine-indifferent to endemic. Serpentine-indifferent plants are equally found on serpentine and non-serpentine soils and serpentine endemic species are only found on serpentine soils.

| Plant species | Plant species code | Soil affinity of population | Soil affinity of species as a whole | Rainfall (cm) |
| --- | --- | --- | --- | --- |
| *Caulanthus amplexicaulis* var *amplex* | AMA | Nonserpentine | Serpentine-Indifferent | 41 to 119 |
| *Streptanthus breweri* | BRE | Serpentine | Serpentine | 51 to 241 |
| *Streptanthus hesperidis* | HES | Serpentine | Serpentine | 81 to 119 |
| *Streptanthus polygaloides* | POL | Serpentine | Serpentine | 48 to 175 |
| *Streptanthus tortuosus* | TORW | Serpentine | Serpentine-Indifferent, primarily nonserpentine | 53 to 424 |
| *Caulanthus crassicaulis* | CRA | Nonserpentine | Nonserpentine | 20 to 43 |
| *Streptanthus diversifolius* | DIV | Nonserpentine | Nonserpentine | 71 to 124 |
| *Streptanthus farnsworthianus* | FAR | Nonserpentine | Nonserpentine | 23 to 41 |

### Study design and rhizoplane soil collection

Plants were a subset from a larger experiment as described previously (Pearse et al. 2020). In summary, seeds from each plant species were collected from natural populations then germinated in a planting tray on a mist bench at the UC Davis Orchard Park greenhouse facility starting in November 2015. From December 2015 to January 2016, seedlings at the cotyledon stage were transplanted into their native soil in containers (Stuewe & Sons, St. Louis, Missouri, USA) in an outdoor lath house at ambient temperature at the Orchard Park facility. The native soils were collected along with the seeds from field populations. The experiment imposed seven watering treatments ranging between 8 to 238 mm/month and a subset of these treatments were used for this study. These levels were chosen to represent a field-relevant range of growing season precipitation based on 30 years of precipitation records experienced by this group of species at our collection sites. Plants were placed in a completely randomized design across the greenhouse.

In June of 2016, rhizoplane soil was collected from flowering plants. A total of 72 samples (3 reps x 3 watering treatments x 8 plant species) were used for this experiment and they included three replicates of eight plant species from three watering treatments, low (16 mm/month = ∼13 mL/week), mid (105 mm/month = ∼87 mL/week), and high (194 mm/month = ∼161 mL/week)(Pearse et al. 2017, 2020).

Rhizoplane soil was collected from roots of individual plants by first shaking the entire root system in a 0.9% (w/v) NaCl solution in a 50-mL centrifuge tube. The roots were transferred into another 50-mL centrifuge tube fill with 20 ml of 0.1% (v/v) Tween80 in 0.9%NaCl solution (Barillot et al. 2013). The tube and solution were shaken on a lateral shaker at 10 spm for 90 min. After shaking, the roots were removed and the tubes containing rhizoplane soil were centrifuged at 200 g for 10 min. The supernatant was discarded, and the pellet used to extract DNA using ZR Soil Microbe DNA MicroPrep following manufacturer’s instructions (Zymo Research, Irvine, CA).

### Library prep and sequencing

From DNA extracts, the V4 region of the 16S SSU rRNA was amplified using primers 515F-806R (515F: 5’ - GTGCCAGCMGCCGCGGTAA - 3’; 806R: 5’ - GGACTACHVGGGTWTCTAAT - 3’) (Caporaso et al. 2010). PCR was carried out in 25 µl reactions including 1 µl genomic DNA, 0.5 µl of each 10 µM primer, 12.5 µl of MyTaq Hot Start Red Mix (Bioline), and 10.5 µl of dH2O. PCR reactions were set up on ice to minimize non-specific amplification and primer dimerization. PCR conditions were: denaturation at 94°C for 2 min; 34 amplification cycles of 30 sec at 94°C, 30 sec at 51°C and 30 sec at 72°C; followed by a 10 sec final extension at 72°C. PCR products were visualized using gel electrophoresis and successful samples cleaned using Carboxyl-modified Sera-Mag Magnetic Speed-beads in a PEG/NaCl buffer (Rohland and Reich 2012).

Cleaned PCR products were quantified using the Qubit HS-DS-DNA kit (Invitrogen, Carlsbad CA), pooled in equimolar concentration and samples were submitted to the Centre for Comparative Genomics and Evolutionary Bioinformatics Integrated Microbiome Resource at Dalhousie University for sequencing using the Illumina MiSeq platform (500 cycles v2 PE250). Raw sequences are available at www.ncbi.nlm.nih.gov/sra/PRJNA613384.

### Bioinformatics

Amplicon sequence variants (ASVs) from 16S rRNA amplicons were identified using DADA2 (v1.7.2) (Callahan et al. 2016a). Briefly, paired-end fastq files were processed by filtering and truncating forward reads at position 280 and reverse reads at position 200. Sequences were dereplicated, merged and error-corrected. Chimeras were removed, and the taxonomy assigned using the SILVA database (v128) (Quast et al. 2012; Yilmaz et al. 2014; Glöckner et al. 2017). A phylogenetic tree based on 16S sequences was estimated using the DECIPHER package (v2.8.1) in R to perform multi-step alignment and phangorn (v2.4.0) to construct the 16S tree (Wright et al. 2006; Schliep 2011). The sequence table, taxonomy, and metadata were combined into a phyloseq object and used for further analysis (phyloseq v1.30.0) (McMurdie and Holmes 2013; Callahan et al. 2016b). Using phyloseq, the mitochondria and chloroplast were removed from samples. Low-abundance samples (<1000 reads) were removed and the count data normalized using relative abundance (McMurdie and Holmes 2014; Callahan et al. 2016b; Weiss et al. 2017).

### Statistical analysis

To visualize the community composition of the microbial communities associated with each plant species across watering treatments, the relative abundance of taxa was visualized by aggregating taxa to the phylum level and filtering out taxa below 2%. To determine if plant species or watering treatment affected microbial diversity in the rhizoplane, Shannon diversity was calculated using the ‘estimate_richness’ function in the phyloseq package (1.30.0) and used as a response variable in ANOVA, as Shannon diversity accounts for both evenness and abundance (Kaisermann et al. 2017; Kim et al. 2017). Watering treatment and plant species were used as predictors and soil affinity was used as a random effect in statistical models. To determine if species and water treatments influenced relative abundance of sequences summed by bacterial genera and ASVs, differential abundance analysis using DESeq2 (1.26.0) was used with watering treatment as a predictor.

To test for differences in multivariate dispersion among rhizoplane communities of species and watering treatment, the ‘betadisper’ function from the vegan (v2.5.3) package was used (Oksanen et al. 2018). To determine if species or watering treatment differed in rhizoplane bacterial composition, we used the ‘adonis’ function from the vegan package with Bray-Curtis dissimilarities as the response variable and plant species and watering treatment as interacting predictors. To visualize the similarity and phylogenetic relationships between groups, principal coordinates analysis (PCoA) were plotted based on Bray-Curtis dissimilarity (Bray and Curtis 1957; Lozupone and Knight 2005).

## RESULTS

After quality filtering and removal of non-target sequences, we recovered 3,603,995 reads (average 50,005 per sample) that were grouped into 10,554 amplicon sequence variants. Sampling curves within samples were saturating, indicating a robust sampling of the microbial diversity associated with individual plants. Most samples were predominated by *Proteobacteria* whereas others hosted mainly *Firmicutes* (Figure 1a).

**Figure 1.**
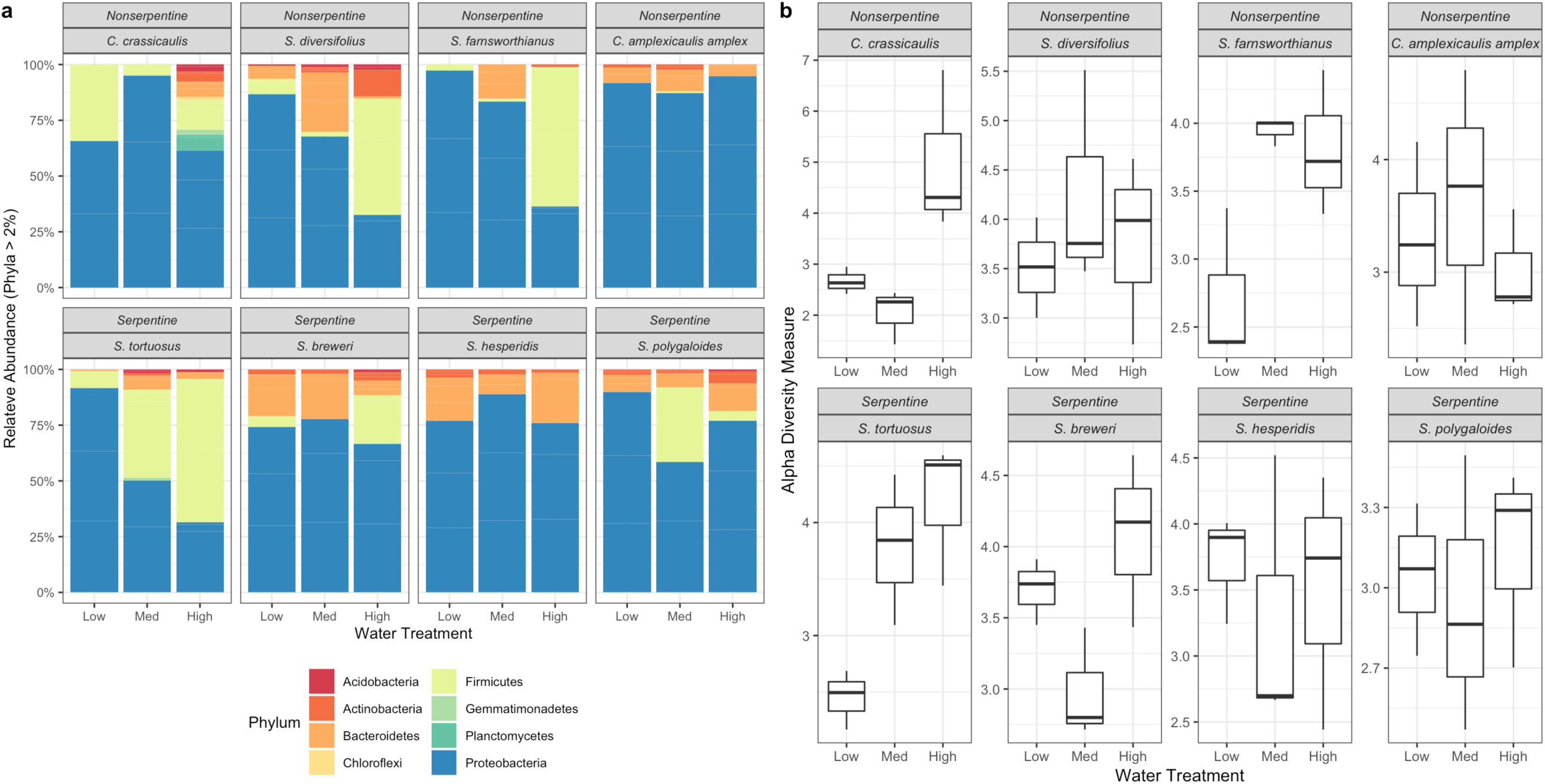
Relative abundance of microbial community. (a) Most samples were predominated by Proteobacteria but Firmicutes were abundant in high water treatments. (b) The alpha diversity between species (Shannon: (F7,48=1.014, P = 0.43) or soil affinity (Shannon: F1,66 = 0.446, P = 0.507) was not significantly different, but it was impacted by watering treatment (Shannon: F2,48 = 5.419, P = 0.008) and the interaction between species and watering treatment (Shannon: F14,48 = 2.734, P = 0.005).

### How does watering treatment impact the alpha diversity of bacteria in the rhizoplane?

Simulated drought significantly decreased Shannon diversity of bacterial communities in the rhizoplane of *Streptanthus* species (Shannon: F_2,48_ = 5.419, *P* = 0.008). Moreover, although plant species did not differ in bacterial alpha diversity (Figure 1b; Shannon: F_7,48_=1.014, *P* = 0.434) watering treatment affected bacterial diversity differently depending on *Streptanthus* species (Shannon: species x watering: F_14,48_ = 2.734, *P* = 0.005). Specifically, alpha diversity was greatest in the high watering treatment for most plant species except for *C. amplexicaulis amplex*, where alpha diversity was the lowest in the high watering treatment.

### How does watering treatment impact the relative abundances of bacterial genera?

Differential abundance analysis using DESeq2 showed that several genera were impacted by watering treatment (Figure 2). The differences in relative abundance were largely driven by the absence of some genera in the low and medium watering treatment. Some phyla had only one genus that significantly changed in relative abundance across watering treatment, whereas others, like Proteobacteria and Firmicutes, had multiple genera impacted by the treatment. Of the Proteobacteria, *H16, Reyranella*, and *Variibacter* increased in relative abundance in the high watering treatment and were absent in the low watering treatment. *Haliangium, Hyphomicrobium*, and *Xanthomonas* were also absent from the low watering treatment, but present in either mid or high watering treatments. Firmicutes including *Psychrobacillus, Sporosarcina*, and *Staphylococcus*, Bacteroidetes such as *Terrimonas* as well as Planctomycetes like *Pirellula* were most abundant in the high watering treatment. DESeq2 analysis of the interaction between watering treatment and species showed that *Flavisolibacter, Paenarthrobacter*, and *Rhodococcus* had different abundances across the watering treatments and species (Supplemental Figure 1).

**Figure 2.**
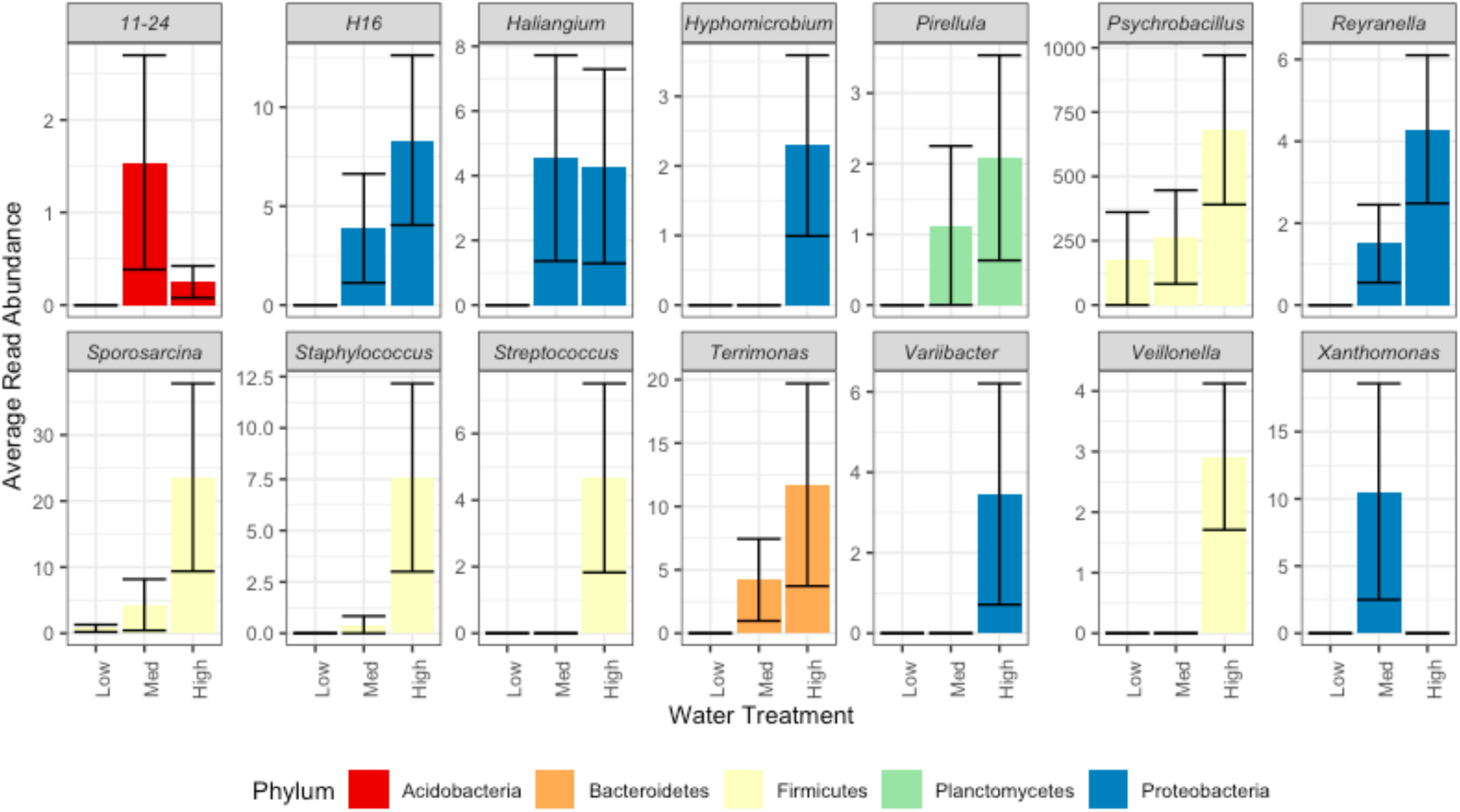
Differential abundances of genera between water treatments using DESeq2. Average read abundance of genera that were shown to be differentially abundant between watering treatments in the rhizoplane at alpha = 0.05. All genera differ significantly between treatments. Bars represent average read abundance with standard error across all plant species.

### How does watering treatment impact the similarity of bacterial communities?

Bacterial species composition differed with watering treatment (Figure 3; F_2,48_ = 1.80; *P* = 0.016) and among plant species (Supplemental Figure 2; F_7,48_ = 1.67; *P* = 0.003). The microbial community variation in the lowest watering treatment was less than the variation in the highest watering treatment (F_2,69_ = 3.80; *P* = 0.032), but there was no significant difference between the variation among plant species (F_7,64_ = 1.71; *P* = 0.125).

**Figure 3.**
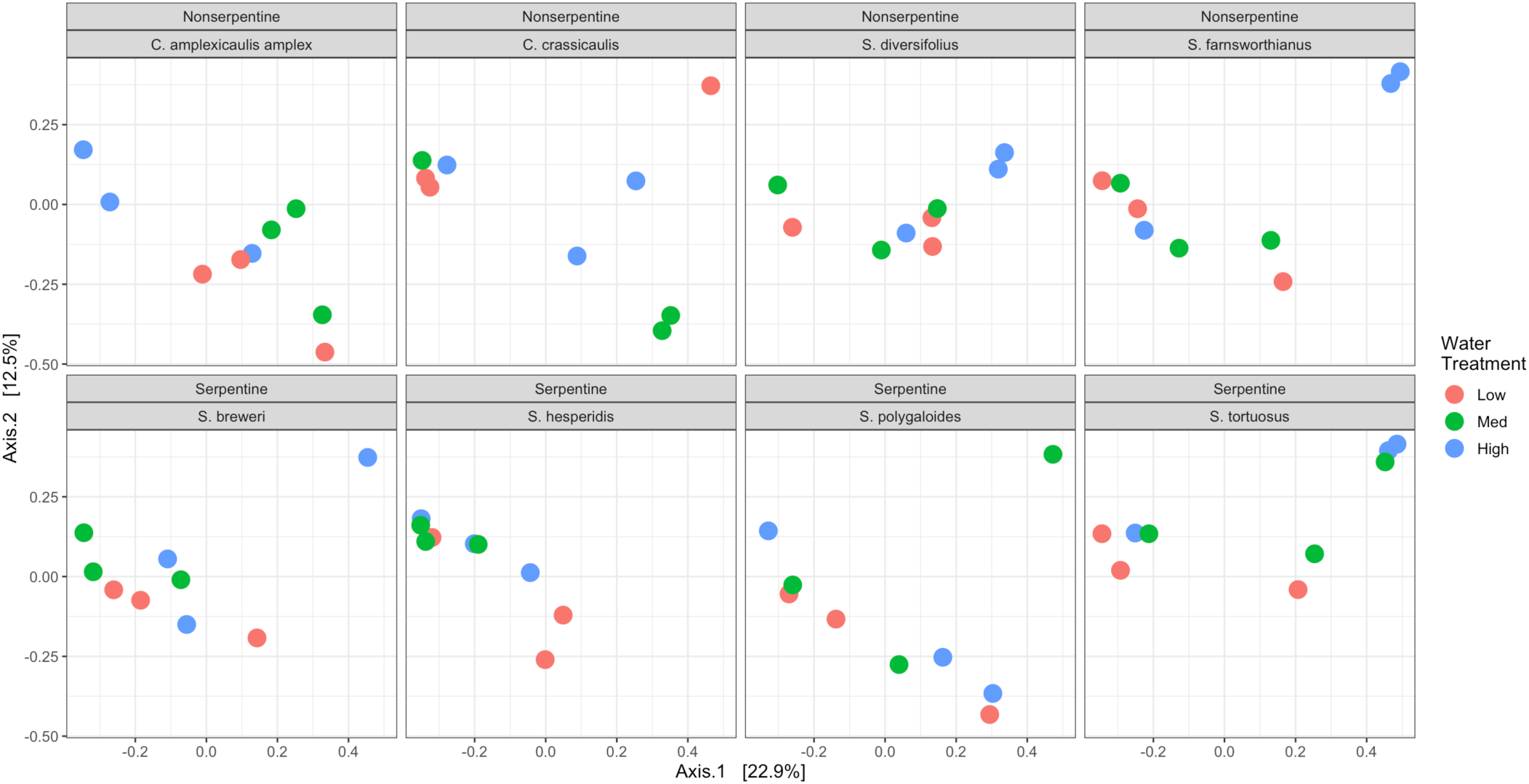
PCoA of rhizoplane samples across watering treatment using Bray-Curtis dissimilarity. Points are rhizoplane microbial communities of associated with *Caulanthus amplexicaulis amplex, Streptanthus brewer*i, *Streptanthus hesperidis, Streptanthus polygaloides, Streptanthus tortuosus, Caulanthus crassicaulis*, S*treptanthus diversifolius*, S*treptanthus farnsworthianus* across watering treatment. Points that are closer together have smaller dissimilarity values meaning they are more similar to each other. Rhizoplane microbial communities become more similar as water stress increases (F2,48 = 1.80; P = 0.016). Points represent individual plants sampled.

## DISCUSSION

### Alpha diversity decreases as water levels decrease

Simulated drought significantly decreased the bacterial diversity and evenness in the rhizoplane across most *Streptanthus* species sampled here. There was a significant interaction between watering treatment and plant species. Unlike most species in the clade, *Caulanthus amplexicaulis amplex*, for example, had the lowest species richness at the highest watering treatment. Although we did not examine the effects of these changes, it may be that changes in microbial diversity either reflect or contribute to variation among *Streptanthus* species to drought conditions (Pearse et al. 2020).

These results mirror previous work, with some studies documenting a reduction in bacterial diversity under water stress (Naylor et al. 2017; Preece et al. 2019; Fahey et al. 2020; Lau and Lennon 2011; Graham et al. 2016; Prudent et al. 2020) while others find little to no effect (Bachar et al. 2010; Tóth et al. 2017). Although we hypothesized that plant soil affinity would predict the effect of watering on microbial communities, our analyses did not support this hypothesis. The variation in the rhizoplane microbial community we observed could have a genetic basis. Members of the same genus often have similar rhizosphere communities (Burns et al. 2015). Measuring the quality and quantity of root exudates of species within the Streptanthus spp. can help us better understand what influences the rhizoplane microbial communities. We did not examine the effect of altered bacterial communities on plant growth or how soil affinity to serpentine soil impacted tolerance to drought. However, in other studies, soil microbial diversity is correlated with improved ecosystem functions (Lau and Lennon 2011; Graham et al. 2016) and in agricultural systems, lower species diversity is often associated with plant yield (Prudent et al. 2020). Although our experimental conditions only approximated drought, the inflection point of fitness (i.e. the amount of water at which adding more water most increased plant fitness) of most of these populations occurred at the watering level reflecting mean annual precipitation for that field collection site (averaged over the last 30 years), suggesting our watering levels are biologically relevant (Pearse et al 2020). This is important because there a different types of droughts which can be characterized based on a lack of precipitation over different time spans as well as increased temperatures and other ecological changes (Hoover and Rogers 2016). Future studies under actual field drought conditions would be useful, particularly if drought duration and watering frequency are important in determining bacterial community responses.

### Plant-Growth-Promoting genera from various phyla disappear in low watering treatments

The relative abundance of several bacterial genera spanning five phyla differed significantly across watering treatments. Genera in the Firmicutes including *Psychrobacillus, Streptococcus, Veillonella, Sporosarcina*, and *Staphylococcus* were more abundant in the highest watering treatment. *Psychrobacillus* is a nitrogen-fixing bacterium often found as a plant endophyte (Rilling et al. 2018; Alishahi et al. 2020) and has been shown, along with *Sporosarcina*, to have plant-growth-promoting properties like phosphorus solubilization and the production of phytohormones (Verma et al. 2016; Xu et al. 2018). In this study, the Bacteroidetes, *Terrimonas*, were abundant in the high watering treatment and they have been shown to be enriched after drought (Meisner et al. 2018; Acosta-Martínez et al. 2014). *Pirellula*, a member of the Planctomycetes, were also the most abundant in the highest watering treatment and they have been implicated as a plant-growth-promoting bacteria in lettuce (Cipriano et al. 2016). In the low watering treatment, these bacterial groups almost disappear from the microbial community. Identifying, isolating, and characterizing drought-adapted microorganisms will be necessary if microbes will be used to assist plant adaptation to climate change.

Many *Streptanthus* grow on rocky outcrops and other stressful environments. Whether specific bacteria contribute to survival or fitness of these plants under stress is not understood. Higher resolution sequencing and functional assays would be useful to confirm if specific bacterial taxa are stress-tolerant and affect plant growth. Further experiments will be required to understand how the loss of these microorganisms under drought will impact the growth and development of *Streptanthus* and *Caulanthus* spp. Some members within the *Streptanthus* genus flower early in the season as a drought avoidance response, and the relationship between drought, accelerated phenology, and the associated microbial community remains to be explored in this genus (Pearse et al. 2020; Igwe et al. 2021).

### Rhizosphere microbial communities become more similar under drought conditions

Our results show that simulated drought shifts rhizoplane microbial communities to be more similar as water levels decrease (Figure 2). Previous research found similar results with C3 and C4 grasses with varying tolerance to water stress (Naylor et al. 2017). In fact, conserved shifts in microbial communities under drought is relatively common (Fitzpatrick et al. 2018; Xu and Coleman-Derr 2019). The finding in this study is somewhat remarkable, given that each species was grown in its own, field–collected soil; soils were of different parent material, and many collection sites were hundreds of miles apart. The affinity to serpentine soil did not impact microbial community composition. It is important to note that although the shift observed in this study is common, the depletion of Gram-positive bacteria is not (Fuchslueger et al. 2014; Schimel et al. 2015). This difference could be due to the initial differences in the bacterial community of field soils. It is worth exploring what mechanisms underlie this trend.

It is important to understand the mechanisms of plant-microbe associations if we are to preserve biodiversity in the face of increasingly variable water regimes, yet these interactions can be context dependent and conditioned by the host plant species (Ishaq et al. 2017; Jorquera et al. 2017; Isobe et al. 2020). Because serpentine soils are naturally drought-prone, we hypothesized that the rhizoplane microbial communities associated with plants with a serpentine-affinity would be less impacted by drought due to legacy effects. However, we did not find support for that hypothesis. Most species in the *Streptanthus* clade grow in well drained, coarse soils (Cacho and Strauss 2014). This may mean that organisms on serpentine soils are not uniquely situated to resist unusual climatic events or that *Streptanthus* species are generally adapted to drought, whether they are associated with serpentine or not. Microbes from drier areas have been shown to use carbon more efficiently than microbes from wetter areas even as their compositions remain the similar under water stress (Leizeaga et al. 2021). Future research would be useful to discern if microbes from serpentine soils use resources more effectively than microbes from nonserpentine soils.

## ACKNOWLEDGEMENTS

I would like to acknowledge Morinne Osborne and Jessica Broaden from the Evolution and Ecology Graduate Admissions Pathways Program at UC Davis for their help collecting samples.

## DATA AVAILABILITY

Data and code available at https://github.com/anigwe/streptanthus_drought

