## Supplemental Figure 1 for "Plant species within Streptanthoid Complex associate with distinct microbial communities that shift to be more similar under drought"

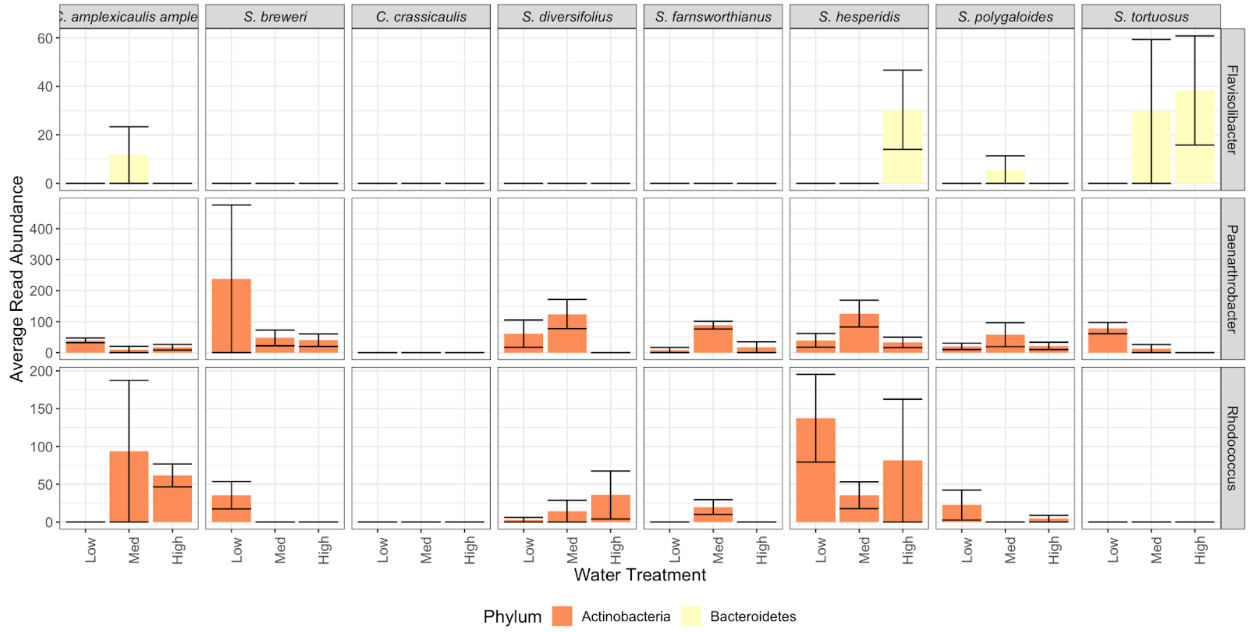


*Supplemental Figure 1 - Differential abundances of genera between species at varying water treatments using DESeq2*

Average read abundance of genera that were shown to be differentially abundant between species at differing water treatments in the rhizoplane at alpha = 0.05. All genera differ significantly between treatments. Bars represent average read abundance with standard error within the plant species and water treatment.
