## Supplemental Figure 2 for "Plant species within Streptanthoid Complex associate with distinct microbial communities that shift to be more similar under drought"

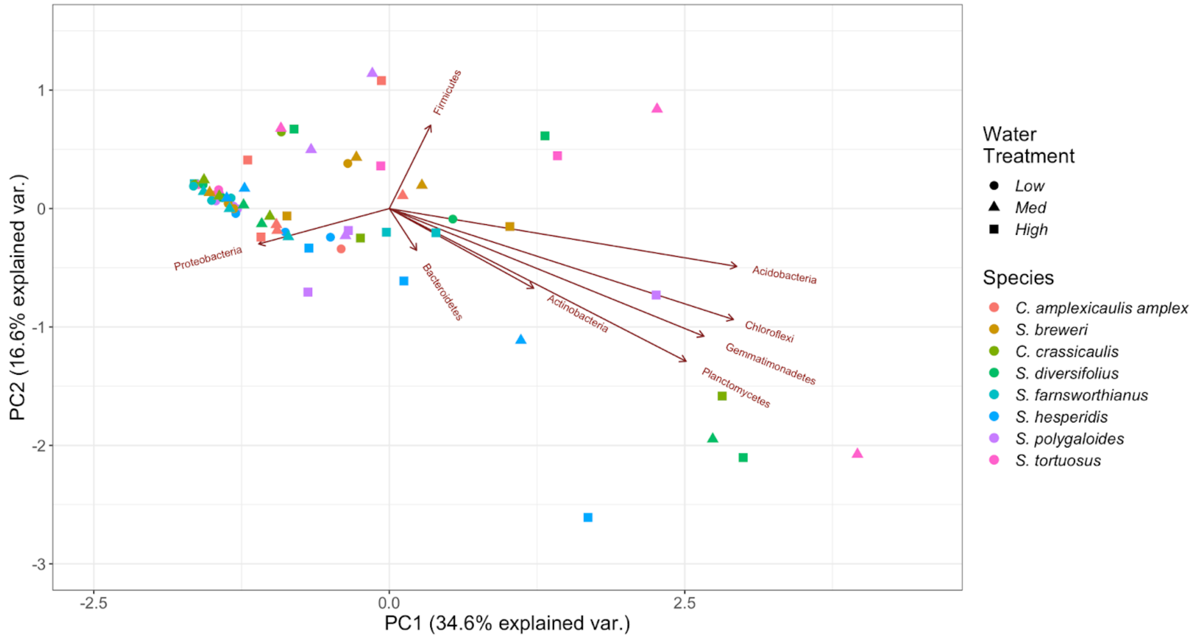


*Supplemental Figure 2 - Principal coordinate analysis (PCA) of the variation rhizosphere microbial communities between species and water treatments*

Each point represents three samples and points that are clustered more closely are more similar. Samples in the low water treatment are clustered tightly at around PC1 = -1 and PC2 = 0. These samples are more similar to each other and Proteobacteria are responsible for much of the similarities. The samples are more spread out in the higher water treatments and are less similar to each other. Members of the Chloroflexi, Actinobacteria, Acidobacteria, Gemmatimonadetes, and Planctomycetes are negatively correlated with Proteobacteria and responsible for much of the dissimilarity in higher water treatments.
